# Coordinating With a Robot Partner Affects Action Monitoring Related Neural Processing

**DOI:** 10.1101/2021.03.26.437133

**Authors:** Artur Czeszumski, Anna L. Gert, Ashima Keshava, Ali Ghadirzadeh, Tilman Kalthoff, Benedikt V. Ehinger, Max Tiessen, Mårten Björkman, Danica Kragic, Peter König

## Abstract

Robots start to play a role in our social landscape, and they are progressively becoming responsive, both physically and socially. It begs the question of how humans react to and interact with robots in a coordinated manner and what the neural underpinnings of such behavior are. This exploratory study aims to understand the differences in human-human and human-robot interactions at a behavioral level and from a neurophysiological perspective. For this purpose, we adapted a collaborative dynamical paradigm from Hwang et al. (1). All 16 participants held two corners of a tablet while collaboratively guiding a ball around a circular track either with another participant or a robot. In irregular intervals, the ball was perturbed outward creating an artificial error in the behavior, which required corrective measures to return to the circular track again. Concurrently, we recorded electroencephalography (EEG). In the behavioral data, we found an increased velocity and positional error of the ball from the track in the human-human condition vs. human-robot condition. For the EEG data, we computed event-related potentials. To explore the temporal and spatial differences in the two conditions, we used time-regression with overlap-control and corrected for multiple-comparisons using Threshold-Free-Cluster Enhancement. We found a significant difference between human and robot partners driven by significant clusters at fronto-central electrodes. The amplitudes were stronger with a robot partner, suggesting a different neural processing. All in all, our exploratory study suggests that coordinating with robots affects action monitoring related processing. In the investigated paradigm, human participants treat errors during human-robot interaction differently from those made during interactions with other humans.

## Introduction

We constantly interact with other humans, animals, and machines in our daily lives. Many everyday activities involve more than one actor at once, and groups of interacting coactors have different size. Especially, interactions between two humans (so-called dyadic interactions) are the most prevalent in social settings (2). During such situations, we spend most of our time trying to coordinate our behavior and actions with other humans. Until recently, human cognition was mostly studied in non-interactive and single participant conditions. However, due to novel conceptual and empirical developments, we are now able to bring dyads instead of single participants to our labs (3). This approach is called Second-person neuroscience (3, 4). It suggests that we need to study the social aspect of our cognition with paradigms that include real-time interactions between participants instead of the passive observation of socially relevant stimuli (4). Such an approach can yield a new perspective on human social cognition.

Coordination between members of a dyad is achieved by joint actions (5). There are different aspects of coordination that facilitate achieving common goals between co-actors. Firstly, (6) showed in pairs of pianists performing solo and duets that monitoring of our actions, our partner’s actions, and our joint actions is required to coordinate successfully. Second, being familiar with each co-actors individual contributions in the dyad helps to form predictions about the partner’s actions, which further improves coordination (7). Third, recently proposed action-based communication serves as a fundamental block of coordination (8). In comparison to verbal communication, this low-level sensorimotor communication is implicit and faster. Experiments by (9) serve as examples of sensori-motor communication in the temporal dimension. Their results have shown that participants adjusted their actions to communicate task-relevant information. Fourth, while both co-actors are engaged in a constant flow of perceptual information, they create coupled predictions about each other’s actions that are necessary to achieve fruitful coordination (5). (10) investigated coordination tasks with incongruent demands between partners, and their results suggested the benefits of reciprocal information flow between participants. In sum, there are different aspects of human cognition that allow for the maintenance of dyadic coordination: Action monitoring, predictions based on familiarity of partner’s actions, action-based communication, and reciprocal information flow.

So far, most dyadic interaction studies investigated the co-ordination between human co-actors (11, 12). However, in recent years we are more and more surrounded by robotic co-actors (13). Furthermore, there are many different predictions for the future of robotics, but all point into the same direction: there will be more robots among us (14). In line with this, humanoid robots are getting progressively better at socially relevant tasks (15). It is thought that these social robots will be used in many different fields of our everyday life in the upcoming years (16). One of the main challenges in robotics is creating robots that can dynamically interact with humans and read human emotions (17). Concerning these changes in our environment, a new research line has emerged and already substantially contributed to our understanding of human-robot interactions (18). As many different scientists are slowly approaching this topic, the field of human-robot interaction until now focused on human thoughts, feelings, and behavior towards the robots (19). Studying these specific aspects is essential and further, we believe that the scientific community has to investigate real-life interactions between humans and robots in order to fully understand the dynamics that underlie this field. Therefore, we propose to use both human and robot partners in experimental paradigms as this will help to close the gap in understanding dyadic interactions.

There are different tools and methods to study the social brain and behavior (20): EEG (21), fMRI (22), MEG (23), and fNIRS (24). From this list, Electroencephalography (EEG) stands out as particularly useful for studying dynamical interactions, as it not only aligns with the temporal resolution of social interactions, but also allows for free movement and thereby allows for dynamic interactions. This temporal resolution allows studying brain processes with milliseconds precision. One of the methods that are classically used within EEG research are event-related potentials (ERPs) (21). ERPs are suitable to study different components of brain processes while they evolve over time. The classic study by (25) showed different brain signatures for correctly and incorrectly performed trials at around 200-300 milliseconds after the feedback about an action was perceived. This brain component was named Feedback related negativity (FRN). In similar studies, (26) showed that the FRN is sensitive not only to our own actions but also those of others. (27) further extended this finding to different social contexts (cooperation and competition). Thus, EEG and specifically ERPs have been proven valuable tools to investigate the physiological basis of social interactions.

However, these studies used static stimuli, and participants performed their actions independently from each other. In the last years, more dynamic experiments and real-life studies are proposed and used within the field of neuroscience (28, 29). Yet, little is known about action monitoring in such dynamic situations with non-human, robotic partners.

To fill this gap, we adapted a dynamic dyadic interaction paradigm for human-robot interactions. We chose the paradigm from (1) and (30), in which two human participants had to manipulate a virtual ball on a circular elliptic target displayed on a tablet and received audio feedback of the ball’s movement. Participants used their fingers to move the tablet and manipulate the position of the ball. We changed the paradigm, by adapting the tablet to enable coordination with the robot and to fit the requirements for EEG measurements. On the one hand, this paradigm allows for coordination similar to a real-life situation; on the other hand, it allows for the analysis of neural underpinnings of cognitive functions required for coordination. In this study, we specifically focused on the aspect of action monitoring with human and robot partners. Thus, to extend our knowledge the present study investigates action monitoring in a dynamic interaction task between humans and robots. Additionally, based on the results from (1) we decided to test whether auditory feedback about actions (sonification) influences coordinated behavior and cognitive processes. Taken together, this study tries to approach a novel problem with interdisciplinary methods and sheds new light on the neural processes involved in dynamic human-robot interactions.

## Methods

### Participants

We recruited 16 participants (7 female, mean age = 25.31 ± 1.92 years) from KTH Stockholm Royal Institute of Technology. We had to exclude two dyads from further analysis, one due to measurement errors in the robot control and one due to excessive movements from participants which led to large artefacts in the EEG data, leaving data from 12 participants in 6 recording sessions. Participants had normal or corrected-to-normal vision and no history of neurological or psychological impairments. They received course credits for their participation in the study. Before each experimental session, subjects gave their informed consent in writing. Once we obtained their informed consent, we briefed them on the experimental setup and task. All instructions and questionnaires were administered to the participants in English. The Swedish Ethical Review Authority (Etikprövningsnämnden) approved the study.

### Task and Apparatus

During each recording session, participants performed the task in four periods of 10 minutes each, twice with a human partner and twice with the robot. Further, each dyad (partner human or robot) performed the task with or without auditory feedback (sonification on or sonification off). The task was based on a tablet game where the dyads cooperated with each other to balance a ball on a circular track as they simultaneously moved it in counter-clockwise direction (1) (Figure 4). At random intervals, we added perturbations that radially dispersed the position of the ball away from the current position. In order to reduce the subjects’ expectations of the occurrence of the perturbations, we sampled its rate of occurrence from a Poisson distribution with *λ*=4s.

The experimental task was implemented on an Apple IPad Air tablet (v2, 2048 x 1536 pixel resolution, refresh rate 60Hz) using Objective-C for iOS. During the task, subjects saw a red ball of 76.8 pixel radius on a circular track with a radius of 256 pixels and a thickness of 42.67 pixels. The ball position was represented as the horizontal and vertical coordinates with respect to the center of the circular track (0,0). The tablet was mounted on a metal frame of size 540mm x 900mm. We further added a square of size 100 pixels x 100 pixels that was used as a signal source for, and covered by, a luminance sensor. Figure 1 shows all the visual components displayed to the participants (the text box on the left side was used by the experimenter to monitor the experiment status). During the periods with another human partner, we asked the participants to not verbally interact with each other. During the task, they sat face-to-face at 1m distance as they held handles connected to the short end of the frame. Similarly, while performing the task with the robot, subjects held the short end of frame while the other end of the frame was clamped to the grip effectors of the robot. Figure 3 shows the physical setup of the subjects and the robot during the experiment.

**Fig. 1.**
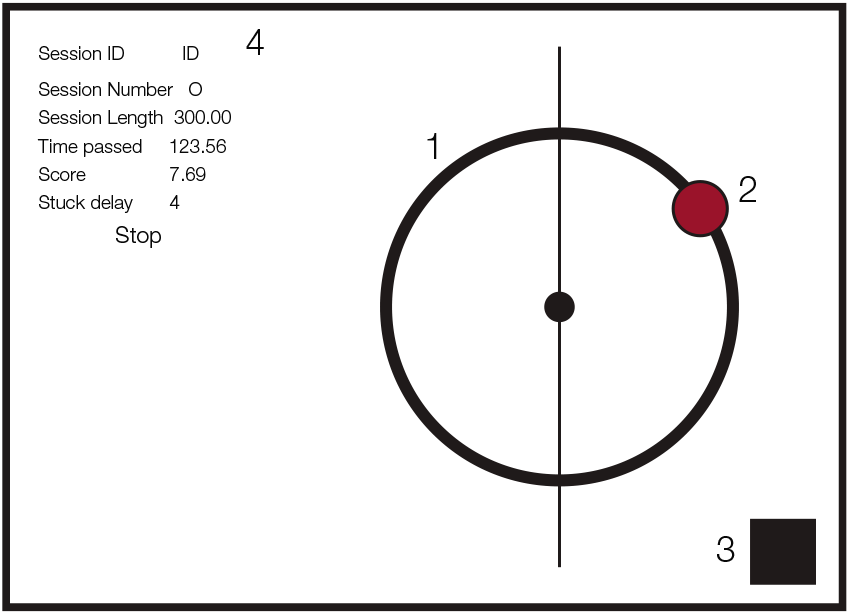
Schematic of game design on the tablet. (1) Circular track, (2) ball, (3) flashing rectangle indicating experimental events (covered by luminance sensor), (4) text box for experiment monitoring (only used by experimenter).

For the periods involving sonification, the position and angular velocity of the ball were sonified. The auditory feedback was created by a Gaussian noise generator with a band-pass filter (cut-off frequency: ± 25Hz). The horizontal and vertical coordinates of the ball modulated the pitch of the auditory feedback, while its angular velocity modulated the loudness. The sonification procedure was implemented using the specifications provided in (1).

Lastly, we used a self-manufactured luminance sensor that synchronized the experimental events (experiment start and end, and perturbation) between the tablet and the EEG amplifier. We changed the luminance source color from black to white to mark the start of the trials, white to black to mark the end of the trials. During a session the patch was white, except at the frame where the perturbation happened, which was marked with grey (RGB=134,134,134).

### Robot Control

We used the YuMi robot (ABB, Västerås, Sweden) shown in Figure 2 for our experiments. We implemented a Cartesian space controller based on the original joint-level velocity controllers provided by the manufacturer. The robot had direct access to the tablet data and no active sensing was necessary. Starting the robot at the joint position depicted in the figure 2, we send Cartesian space velocity commands to both arms at 10 Hz. The Cartesian controller was designed such that the *X, Y* positions of both end-effectors are kept constant during an execution, and only the *Z* position of the end-effectors are adjusted to move the ball. We denote the left and right end-effector velocity commands in the *z* axis by 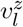 and 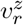 and the current *X, Y* position of the ball on the game by (*b*_*x*_, *b*_*y*_), respectively. We first obtain the angle *θ* corresponding to the current position of the ball in the polar coordinate system by *θ* = *arctan*(*b*_*y*_, *b*_*x*_). Then, we obtain the next target angle 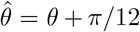 to let the ball move in the counterclockwise direction. The next target *X* ,*Y* positions of the ball are found as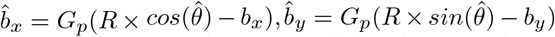, where *R* is the radius of the circle on the IPad game and *G*_*p*_ = 0.1 is a constant gain. The velocity commands in the *z* axis are then found as 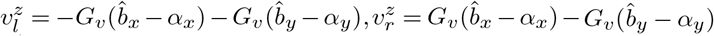, where, *α*_*x*_, *α*_*y*_ are gravity acceleration in the *X, Y* directions measured by the IPad, and *G*_*ν*_ = 0.5 is a constant gain. The command velocities are then clipped to have an absolute value less than 0.02 m/s, and the clipped values are sent to the Cartesian velocity controller.

**Fig. 2.**
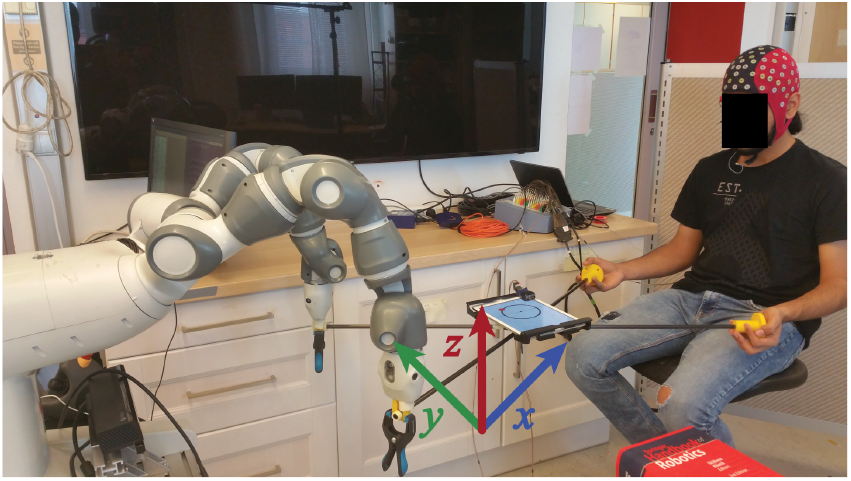
Robot arms and their degrees of freedom. The participants played with a Yumi robot. Its arms were able to move the tablet in 3D space (x, y, z).

**Fig. 3.**
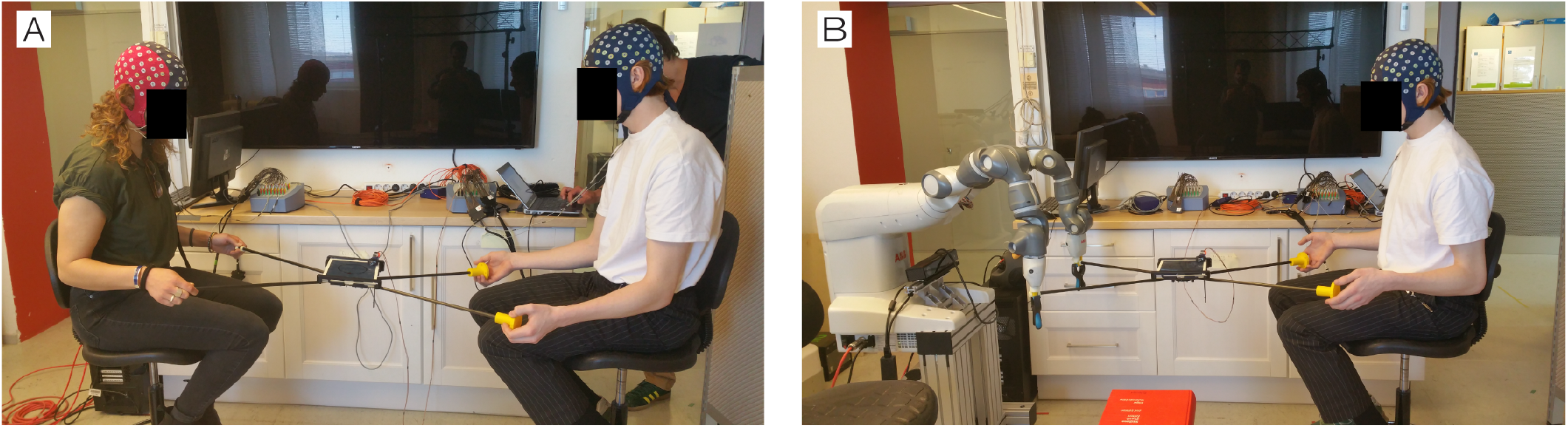
Experimental Setup. Participants performed the experiment with a human (A) or a robot partner (B). In each condition they played a tablet game by balancing a virtual ball on a circular track while moving it in the counter-clockwise direction.

### Procedure

We prepared both participants for the EEG recording together, which took around 45 minutes to complete. Once the subjects were ready to start the experiment, we led them to a room that housed the robot. Depending on the dyad combination, we provided oral instructions about the task and clarified any remaining questions. For the human-robot dyads, we first reset the limbs of the robot to its initial conditions and then started the task on the tablet. After each block, the participants were given a short break and then repeated the task with the alternate sonification condition. The whole experimental session lasted for about 4 hours.

### EEG data acquisition

We recorded the EEG using two 64-Ag/AgCl electrode systems (ANT Neuro, Enschede, Netherlands), and two REFA8 amplifiers (TMSi, Enschede, Netherlands) at a sampling rate of 1024 Hz. The EEG cap consists of 64 electrodes placed according to the extended international 10/20 system (Waveguard, eemagine, Berlin, Germany). We placed the ground electrode on the collar-bone. We manually adjusted the impedance of each electrode to be below 10kΩ before each session. The recording reference was the average reference, which, only in the single-brain recordings, was later programatically re-referenced to Cz. During human-human interactions, two brains were recorded simultaneously with the separate amplifiers, synchronized through the ANT-link (Synfi, TMSi, Enschede, Netherlands). VEOGs were recorded with two additional electrodes, one placed below and one above the eye.

### Pre-processing

The analysis of the EEG data was performed in MATLAB 2016b and the behavioral analyses in Python 3.7.

We preprocessed the data using the EEGLAB toolbox (v2019.0) (31). As a first step before preprocessing, we programmatically extracted the trigger events from the luminance sensor and added them to the recorded data. Then, the data from each condition was downsampled to 512Hz, followed by referencing all datasets to Cz electrode. We then high-pass filtered the dataset at 0.1Hz and then low-pass filtered it at 120Hz (6 dB cutoff at 0.5Hz, 1 Hz transition band-width, FIRFILT, EEGLAB plugin, (32)). Following this, we manually removed channels that showed strong drift behavior or excessive noise (mean: 7, SD: 2.7, range: 1-13). We manually inspected the continuous data stream and rejected the portions which exhibited strong muscle artifacts or jumps. To remove further noise from eye and muscle movements, we used independent component analysis (ICA) based on the AMICA algorithm (33). Before performing ICA, we applied a high-pass filter to the data at 2Hz cut-off to improve the ICA decomposition (34). We visually inspected the resulting components in combination with using ICLabel (35) classifier. IClabel was run on epoched data, 200 ms before and 500 ms after the perturbation. Based on the categorization provided by ICLabel, and a visual inspection of the time course, spectra, and topography, we marked ICs corresponding to eye, heart and muscle movements for rejection (mean: 26.5, SD: 5.2, range: 18-44). We copied the ICA decomposition weights to the cleaned, continuous data and rejected the artifactual components. Finally, using spherical interpolation, we interpolated the missing channels based on activity recorded from the neighboring channels.

### Behavioral Analysis

In order to understand the behavioral differences for the factors partner and sonification, we used measures of mean angular velocity and mean error produces. We first calculated the instantaneous angular position *θ* (in degrees) of the ball using the horizontal and vertical (X, Y) positions of the ball on the tablet as follows:

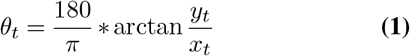

We used the atan2 function to take into account the X, Y position in the negative coordinate axes. *θ*_*t*_ values were transformed from [−*π, π*] to range [0, 2*π*]. Next, we computed the instantaneous angular velocity *ω* of the ball using the following formula where *t* is the sample time-point:

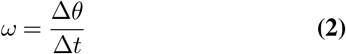

We, subsequently, calculated the mean *ω* for each participant for the four different conditions. Next, We calculated the error as the difference of the instantaneous radial distance between the radius of the track and the ball’s current position measured as the distance from the track’s center as follows:

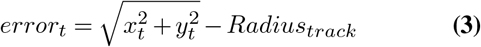

### Deconvolution and EEG Analysis

Even though the perturbations were sampled from a Poisson distribution with *λ* = 4, the corresponding neural responses might overlap in time and bias the evoked potentials. Further, block onset and offset typically elicit very strong ERPs overlapping with the perturbations. Finally, we see clear, systematic differences in the behavior depending on the condition (e.g. higher velocity with a human partner), which could lead to spurious effects in the ERPs. We further added eccentricity (distance from the circles midpoint), in order to control for the ball’s trajectory. In order to control both temporal overlap and covariate confounds, we used linear deconvolution based on time-regression as implemented in the unfold toolbox v1.0 (36). Consequently, we modeled the effects of the partner (human or robot), the sonification (off = 0, on = 1) and their interaction as binary, categorical variables, the eccentricity and the velocity were coded using B-spline basis functions and the angular position using a set of circular B-splines. The block on- and offsets were modeled as intercept only models. The complete model can be described by the Wilkinson notation below (37).

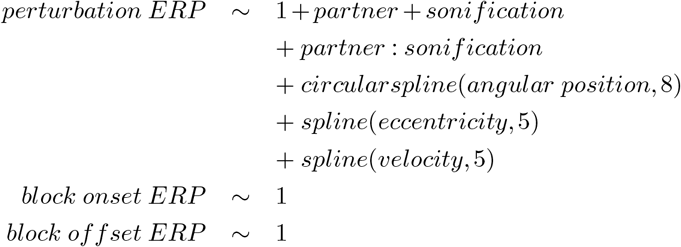

This model was applied on the average referenced continuous EEG data, and each event was modeled in the time range of -500ms to 700ms with respect to the event onset.

Similar to the two-stage mass univariate approach, we calculated the t-value over subjects for each of the resulting regression coefficients (similar to difference waves) for all electrodes and time points (time-range of -500ms to 700ms). The multiple comparison problem was corrected using a permutation based test with threshold-free cluster enhancement (TFCE) (38, 39) with 10,000 permutations(default parameters *E* = 0.5 and *H* = 2). We used the eegvis toolbox (40) to visualize all evoked response potentials.

## Results

### Behavioral

In this study, humans played a collaborative game either with other humans or with robots. We further added sonification of the ball’s movement as a supplementary auditory feedback to the participants. Figure 4 shows the raw positions of the ball overlaid for all subjects and the partner and sonification conditions. The behavior we analyse here, is the mean velocity of the ball during each session and the mean deviation of the ball from the circular track. These measures indicate how fast the participants performed the task and how much error they produced, both a proxy of the success of the collaboration.

**Fig. 4.**
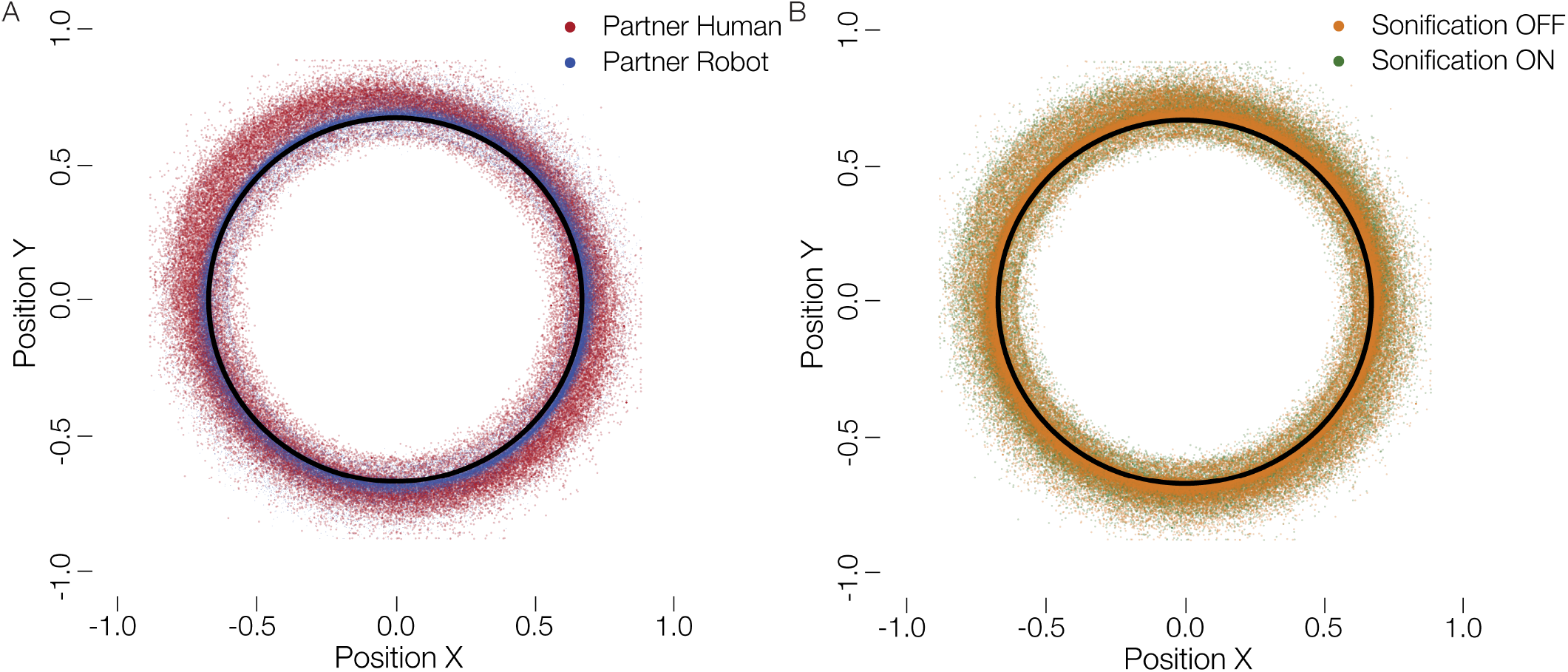
Distribution of ball positions. (A) Ball positions on the tablet with a human (red) and robot (blue) partner. (B) Ball positions on the tablet for sonified (green) and not sonified movements (orange). The black circle represents optimal trajectory. It can be seen that participants deviated more with a human partner. No such difference is visible for a change in the sonification.

We calculated the mean angular velocity (*ω*) for each participant for the four different conditions (Figure 5 (A)). To test the statistical significance of these findings, we computed a 2×2 factorial repeated measures ANOVA with the factors partner and sonification. The ANOVA showed a significant main effect of partner, *F* (1, 11) = 87.09, *p <* .0001 where subjects exhibited a mean angular velocity of 265.20 degrees/second and SD ± 0.28.29 with a human partner, conversely, with a robot partner subjects showed a mean angular velocity of 159.23 degrees/second ± 29.40. The ANOVA did not yield a significant main effect of sonification, *F* (1, 11) = 1.00, *p* = 0.33, with mean angular velocity 210.06 degrees/second ± 65.51 with sonification off and the mean angular velocity was 214.36 degrees/second ± 62.53 with sonification on. There was no significant interaction of factors partner and sonification, *F* (1, 11) = 0.04, *p* = 0.83. Hence, we can conclude that participants were faster at moving the ball on the circular track while performing the task with a human partner.

**Fig. 5.**
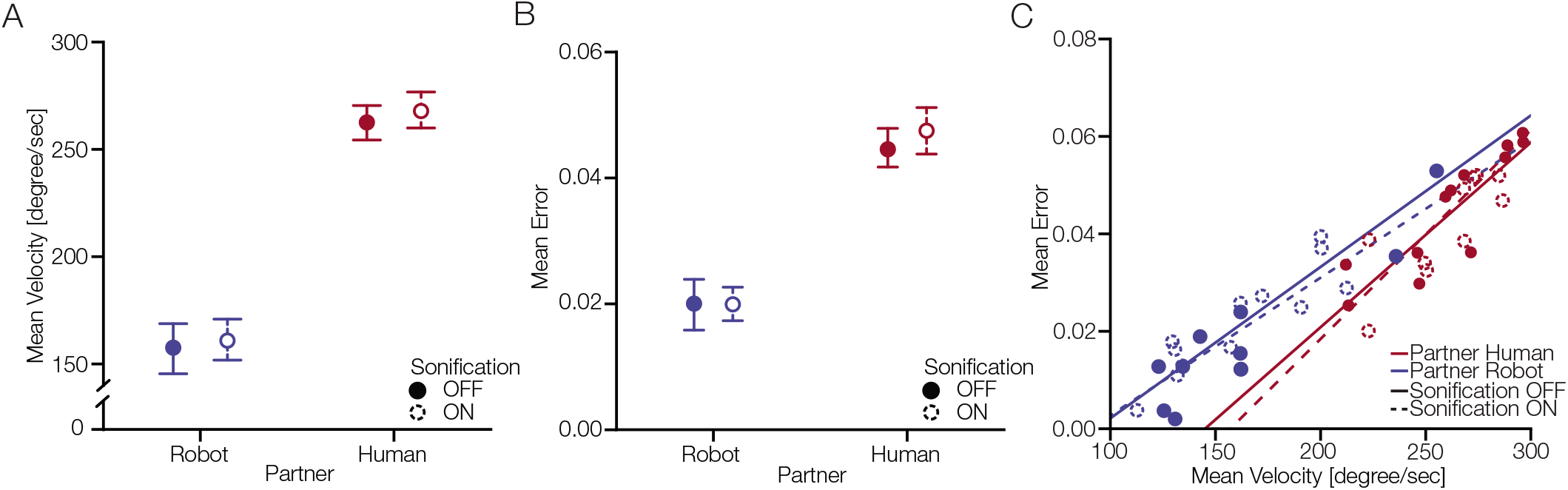
Behavioral differences between conditions partner and sonification. (A) Differences in mean angular velocity across different participants. The error bars indicate standard error of mean. (B) Differences in mean error across different participants. The error bars indicate standard error of mean. (C) shows the correlation of mean error and mean velocity for partner and sonification conditions.

Next, we analysed the mean error produced by participants during a session. Figure 5 (B) shows the mean error across participants for the four different conditions. To statistically assess these differences, we performed a 2×2 factorial repeated measures ANOVA with factors partner and sonification. The ANOVA yielded significant main effect for partner *F* (1, 11) = 42.61, *p <* .0001 where subjects had a mean error of 0.04 ± *SD* = 0.012 while performing with a human partner, conversely, they had a mean error of 0.01 ± 0.012 while cooperating with the robot. We did not find a significant main effect of sonification *F* (1, 11) = 1.75, *p* = 0.21 where subjects had a mean error of 0.032 ± 0.017 with the sonification off and mean error of 0.033 ± 0.018 with sonification on. There was no significant interaction of factors partner and sonification, *F* (1, 11) = 0.51, *p* = 0.48. We can conclude that subjects made larger errors while performing the task with a human partner compared to the robot partner. Lastly, we were interested in the correlation between the behavioral measures we analysed. Figure 5 (C) shows the correlation of mean error and mean velocity for the partner and sonification conditions. For human partner with sonification off the Pearson correlation showed a correlation coefficient *ρ* = 0.98, *p <* 0.001 and for sonification on *ρ* = 0.89, *p <* 0.001. For robot partner with sonification off *ρ* = 0.97, *p <* 0.001 and with sonification on *ρ* = 0.97, *p <* 0.001. These results show that the mean error and mean velocity were positively correlated during the task.

### EEG

Next, we look at the brain activity during the task. Using a overlap-corrected time regression approach, we investigate the main effect and interaction ERPs from the 2×2 design, while adjusting for eccentricity, velocity and position of the ball (see Methods for details). We only report ERPs time-locked to perturbation events.

Descriptively, in electrode Cz (Figure 6 (A)), we see the typical pattern of a positive inflection, followed by a negative and a second positive inflection after the perturbation onset. We did not have a specific hypothesis to a predefined component and analyzed all electrodes and time points simultaneously. The TFCE analysis reveals two clusters for the main effect of the factor of partner (Figure 6 (B)). The first cluster is likely to represent the activity between 230ms and 270ms with its maximum amplitude being -2.8µV at electrode FC1 (median p: 0.025, minimal p: 0.018). The second cluster most likely represents the time range of 515ms to 605ms with a peak at - 1.2µV at electrode FC2 (median p: 0.026, minimal p: 0.002). Both clusters are found in the central region. No significant clusters were found for neither the factor sonification nor the interaction term.

**Fig. 6.**
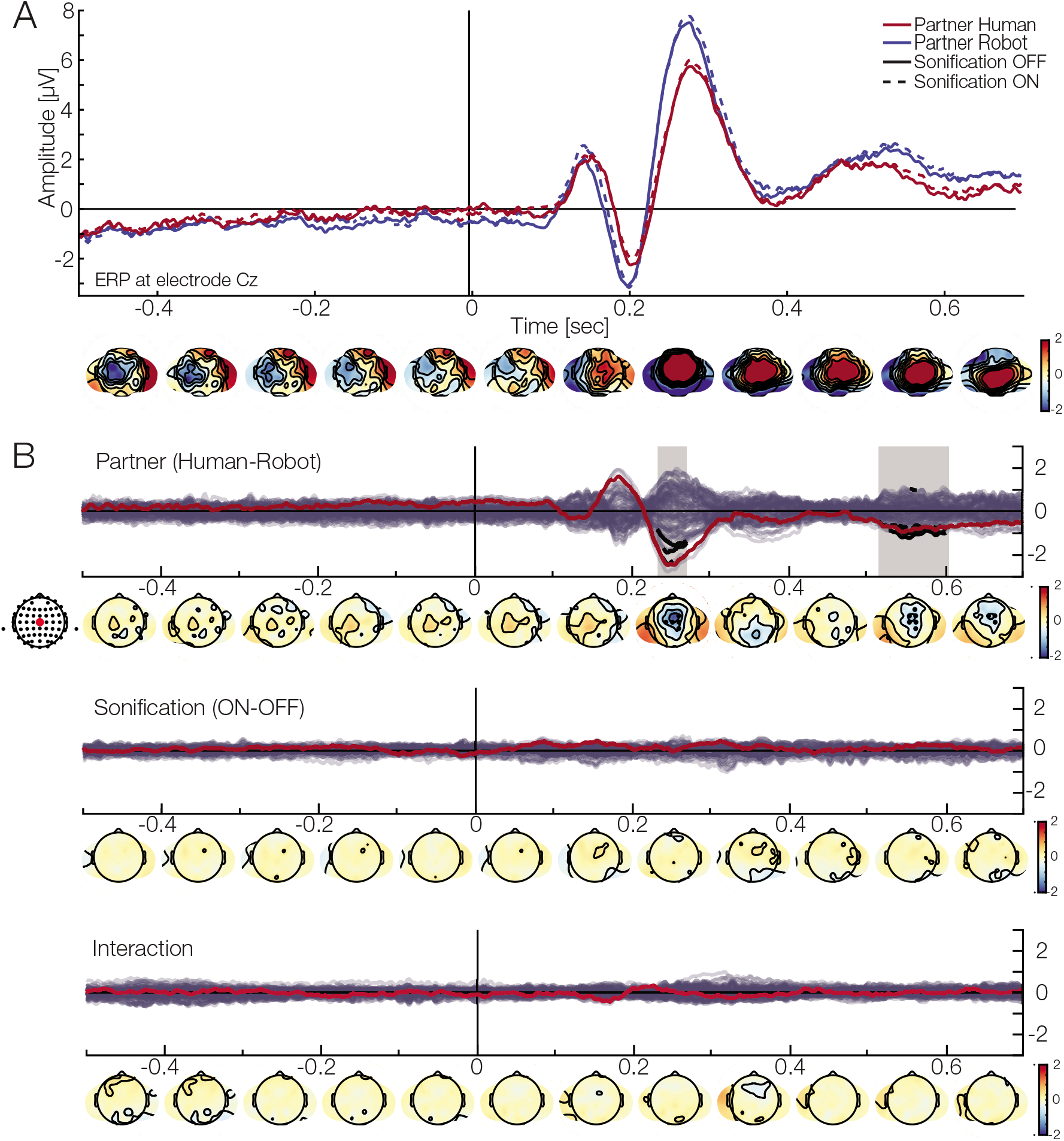
EEG results. (A) ERP at electrode Cz. The red lines show the activation when interacting with a human partner, while blue lines indicate a robot partner. The solid line are the ERP when the sonification was off, while the dashed line represent sonification on. Below, are the topographies for the grand average (mean over all conditions). (B) Clustering results for differences between (red line and dot represents electrode Cz). Top: Effect of partner. The analysis finds two clusters in the central area (black dots and segments). One is likely due to a difference at around 230ms to 270ms, while the second one is present later (around 510ms to 600ms). These results indicate that the ERP will have a smaller amplitude when interacting with a human partner. Middle: Effect of sonification. No cluster was found here. Bottom: Interaction. No cluster was found here.

These results show that we find differences in the participants’ ERPs with respect to their current partner independently of their differences in behavior: When interacting with a robot partner the ERP will have a stronger amplitude indicating a systematically different processing.

## Discussion

Dynamic human-robot interaction studies are at the frontiers of joint-action research. Our experiment investigated neural correlates of action monitoring in a dynamic collaboration task that involves two co-actors. Participants performed the task with another human and robot partner while we measured EEG signals. Co-actors tried to keep a virtual ball on the circle displayed on a tablet; they used their hands (human arm or robotic arm) to manipulate independent orientation axes of the tablet. We perturbed the ball to investigate neural action monitoring processes of the participants. We found fronto-central ERP components at around 200-300ms after the ball was perturbed. The components were stronger for human and robot partner compared to interactions with another human. These results suggest that the dynamic processing of our actions is influenced by whether we collaborate with a robot or a human.

The exploratory aspect of investigating neural underpinnings of human-robot interactions pose many challenges and questions. In the present study, we tried our best to reconcile all of them. However, there are limitations that have to be addressed. First, our sample size was small in terms of number of dyads. However, it was not small in terms of recordings and total amount of gathered data. Thus, the effects reported are significant at high level. Second we did not perform statistics with a predefined hypothesis. Instead, we performed an exploratory analysis that encompasses all electrodes and time points. It is important to understand that it is the first study of its kind. Therefore, it has to be replicated and evaluated by future research (41). Third our results could be dependent on the robot used in the study. We suggest that different types of robots (less/more humanoid) could modulate action monitoring differently. The robot used in the present study was clearly not-humanoid. Participants could clearly recognize it as a robot and devoid of typical human traits that are often used in communication/collaboration. Nonetheless, using this robot helped us to maximize the difference between conditions. Additionally, our claim is supported by research on a different level of trust depending on the appearance of humanoid robots (42, 43). Therefore, it would interesting to perform a similar experiment and compare the results with a more human-like robot. Fourth, as discussed below, our robot did not have a model of the human actor. By this, the robot’s behavior helped to boost the characteristic differences between the player’s partners. Fifth, our statistical analysis does not take the dyadic dependency into account, possibly biasing the estimated model parameters of the human-human condition downward. In the future, study with a bigger sample size, could answer the question whether dyadic dependencies play a role in the effects reported in our study. Sixth, even though participants were asked to keep their eyes on the center of the circular track, we did not control for eye-movements in this study, which could result in biased viewing-behavior on the tablet. However, we adjusted for ball position while modeling the ERPs, which is likely to be a proxy for current eye position and also remove eye movement and blink related ICs. Furthermore, the game required constant attention and engagement, so it was assured that participants did not look away from the tablet and the ball. Additionally, we are interested in the EEG signal related to the behavior, rather than the visual stimulus. All in all, we addressed the limitations, and are convinced that they do not impede the interpretations of our results as presented in next paragraphs.

The behavioral measures of our participants’ actions were different between human and robot partners. We focused our analysis on two aspects of collaboration: The speed which is represented by the ball’s velocity and the accuracy as indicated by the mean error. Our results suggest that participants perform slower when paired with the robot and achieve higher accuracy (ball closer to the circular track). There is a trade-off relation between these factors; this is why we discuss them together (Figure 5 (C)). One simple explanation could be that the robot’s control were themselves slow and prone to error. The human participants might have restrained themselves and thereby executed artificially slow movements. Another interpretation of why our participants slow down (and increased accuracy) while performing with the robot is that they had less trust in the robot than a human partner. This is in line with past research that suggests that level of trust changes during real-time interactions with robots (44) and that, in general, trust levels are different for human and robot partners (45). Another interpretation for slower movements is that it is challenging to create a model of a partner’s actions during a joint collaborative task with a robot. Because we typically represent others’ actions as our own (46), it is possible that in the case of interacting with a robot we need more time to create such representations. There is much space for interpretations why having a robot partner triggered slower movements; however, we would like to point that the main goal of our study was to investigate neural correlates of different partners, and behavioral responses were collected to exclude their influence on neural responses (see Deconvolution and Analysis for details).

After adjusting for behavioral differences in the EEG analysis, we see that robot partners affect neural correlates of action monitoring differently in comparison to a human partner. We found that between 200-300 ms after the perturbation event disturbing the collaboration, the EEG amplitudes differ at the fronto-central sites. Concerning exploratory aspect of this study and used analysis, it is challenging to map and compare our results with well established ERP components. The literature on single participants at these electrodes and time window suggests that it is when and where monitoring our errors or feedback about our actions unravels (25, 47). Similarly, when it comes to neural activity involved in action monitoring in dyadic situations, the same activations play a role (26, 27). If the error is committed by the participant and can be inferred from his action (e.g. making a typo), the brain component involved is called Error-related negativity, with more negative activation for erroneous actions than correct ones (48). In case of behavior that needs feedback to understand the consequences of the action (for example, gambling task), it is called Feedback related negativity (49). In comparison to these classic, static, and passive experiments, we had real-time collaboration between two participants, and we observed similar component peaking around 200-300 ms after the perturbation happened. Our participants were not informed about the perturbations, so they could have been treated as participants’ own or the partner’s error. Therefore, we suggest that the neural activation we observe in our study resembles classic components. Importantly, our results do not only resemble previous research but contribute with a new finding: robot partners modulate action monitoring. This result is in line with the other EEG study comparing humans with robot partners (50). There is a crucial difference between both studies: Participants in (50) study performed a task sequentially (turn-taking), while in our study, participants interacted with each other in real-life. Both studies point in the same direction. Robot partners modulate neural activity.

Our results suggest that robot partners can modulate neural activity in a dyadic experiment. Our and (50) studies contribute to our understanding of human cognition. Concerning that there is not many studies that focused on neural underpinnings of human-robot interactions, the results we present here have a value for research topics in the field of join-action. They are a first exploratory step towards a theoretical and methodological foundation. We showed the feasibility of conducting a human-robot interaction study while measuring EEG from the human participant. With full experimental control, we explored neural correlations of human-robot interactions in an ecologically valid setup (29, 51). There is vast literature on the topic of joint actions between humans and robot partners (52, 53). However, what was missing until now, are neural indicators human-robot joint actions. Our study shows that it is possible to conduct studies with non-human agents collaborating with humans and measure brain activity and that the neural basis of action monitoring is affected by the robot partner.

Lastly, we observed small differences between human and robot partners at later time points (between 500-600 ms after the perturbation) around the midline electrodes. These differences are difficult to interpret. The topography suggest similar source as the component discussed above. However, based on time we speculate it could be P3b component. (54) reported similar component in study that investigated self and other (human versus computer) generated actions in pianists. They found that P3b component was present only for self generated actions, suggesting greater monitoring of self generated actions. It is important to highlight that in our study, participant had to dynamically perform the task, while in the (54) study participants took turns to perform joint actions. What is similar is that they had to generate actions to achieve a common joint goal (12). It is possible that the late effect we found in our experiment has the same function (greater monitoring of self generated actions). However, in comparison to the earlier effect (200-300 ms after the perturbation), the size of the effect in our study is small. Therefore, we have to be careful with interpretations. Future researcher with bigger sample size can help to understand the function of late ERP components in joint actions with robots.

Taken together, this study explored and described event-related potentials related to action monitoring in humans collaborating with other humans or robots. We used a dynamic real-time collaborative task and found that around 200-300 ms after our actions are disturbed, our brain activity is modulated by the type of a partner. Our study shows the feasibility of conducting research on collaboration between human and non human partners with EEG. Furthermore, results of our study suggest that non-human partners modulate how we perceive and evaluate joint actions.

## Author Contributions

PK, DK, MB: conceived the study. AC, ALG, AK, PK: designed the study. AG, MB: programmed the tablet and the robot. AC, ALG, AK, AG, MT: data collection. ALG, AK: major data analysis. AC, TK, BE, PK: minor data analysis. AC, ALG, AK: initial draft of the manuscript. AC, ALG, AK, AG, BE, MB, PK: revision and finalizing the manuscript. All authors contributed to the article and approved the submitted version.

## Acknowledgments

We would like to thank all partners in the socSMC consortium. Especially, we thank Alfred O. Effenberg and Tong-Hun Hwang for providing the sonification algorithm, and the help with implementing it.

## Bibliography

1. Tong-Hun Hwang, Gerd Schmitz, Kevin Klemmt, Lukas Brinkop, Shashank Ghai, Mircea Stoica, Alexander Maye, Holger Blume, and Alfred O. Effenberg. Effect-and Performance-Based Auditory Feedback on Interpersonal Coordination. Frontiers in Psychology, 9:404, March 2018. ISSN 1664-1078. doi: 10.3389/fpsyg.2018.00404.

2. Leonard S Peperkoorn, D Vaughn Becker, Daniel Balliet, Simon Columbus, and Catherine Molho. The prevalence of dyads in social life. PLOS ONE, page 17, 2020.

3. Leonhard Schilbach, Bert Timmermans, Vasudevi Reddy, Alan Costall, Gary Bente, Tobias Schlicht, and Kai Vogeley. Toward a second-person neuroscience. Behavioral and Brain Sciences, 36(4):393–414, August 2013. ISSN 0140-525X, 1469-1825. doi: 10.1017/S0140525X12000660.

4. Elizabeth Redcay and Leonhard Schilbach. Using second-person neuroscience to elucidate the mechanisms of social interaction. Nature Reviews Neuroscience, 20(8):495–505, August 2019. ISSN 1471-003X, 1471-0048. doi: 10.1038/s41583-019-0179-4.

5. Natalie Sebanz and Günther Knoblich. Progress in Joint-Action Research. Current Directions in Psychological Science, page 096372142098442, January 2021. ISSN 0963-7214, 1467-8721. doi: 10.1177/0963721420984425.

6. Janeen D. Loehr, Dimitrios Kourtis, Cordula Vesper, Natalie Sebanz, and Günther Knoblich. Monitoring Individual and Joint Action Outcomes in Duet Music Performance. Journal of Cognitive Neuroscience, 25(7):1049–1061, July 2013. ISSN 0898-929X, 1530-8898. doi: 10.1162/jocn_a_00388.

7. Thomas Wolf, Natalie Sebanz, and Günther Knoblich. Joint action coordination in expert-novice pairs: Can experts predict novices’ suboptimal timing? Cognition, 178:103–108, September 2018. ISSN 00100277. doi: 10.1016/j.cognition.2018.05.012.

8. Giovanni Pezzulo, Francesco Donnarumma, and Haris Dindo. Human Sensorimotor Communication: A Theory of Signaling in Online Social Interactions. PLoS ONE, 8(11):e79876, November 2013. ISSN 1932-6203. doi: 10.1371/journal.pone.0079876.

9. Cordula Vesper, Laura Schmitz, and Günther Knoblich. Modulating action duration to establish nonconventional communication. Journal of Experimental Psychology: General, 146 (12):1722–1737, December 2017. ISSN 1939-2222, 0096-3445. doi: 10.1037/xge0000379.

10. Arianna Curioni, Cordula Vesper, Günther Knoblich, and Natalie Sebanz. Reciprocal information flow and role distribution support joint action coordination. Cognition, 187:21–31, 2019.

11. N Sebanz, H Bekkering, and G Knoblich. Joint action: Bodies and minds moving together. Trends in Cognitive Sciences, 10(2):70–76, February 2006. ISSN 13646613. doi: 10.1016/j.tics.2005.12.009.

12. Cordula Vesper, Stephen Butterfill, Günther Knoblich, and Natalie Sebanz. A minimal architecture for joint action. Neural Networks, 23(8-9):998–1003, October 2010. ISSN 08936080. doi: 10.1016/j.neunet.2010.06.002.

13. Mordechai Ben-Ari and Francesco Mondada. Robots and Their Applications, pages 1– 20. Springer International Publishing, Cham, 2018. ISBN 978-3-319-62532-4 978-3-319-62533-1. doi: 10.1007/978-3-319-62533-1_1.

14. Peter Stone, Rodney Brooks, Erik Brynjolfsson, Ryan Calo, Oren Etzioni, Greg Hager, Julia Hirschberg, Shivaram Kalyanakrishnan, Ece Kamar, Sarit Kraus, et al. Artificial Intelligence and Life in 2030. One hundred year study on artificial intelligence: Report of the 2015-2016 Study Panel. Stanford University, Stanford, CA, http://ai100.stanford.edu/2016-report. xAccessed: September, 6:2016, 2016.

15. Riccardo Campa. The rise of social robots: A review of the recent literature. Journal of Evolution and Technology, 26(1), 2016.

16. Sibylle Enz, Martin Diruf, Caroline Spielhagen, Carsten Zoll, and Patricia A. Vargas. The Social Role of Robots in the Future—Explorative Measurement of Hopes and Fears. International Journal of Social Robotics, 3(3):263–271, August 2011. ISSN 1875-4791, 1875-4805. doi: 10.1007/s12369-011-0094-y.

17. Guang-Zhong Yang, Jim Bellingham, Pierre E. Dupont, Peer Fischer, Luciano Floridi, Robert Full, Neil Jacobstein, Vijay Kumar, Marcia McNutt, Robert Merrifield, Bradley J. Nelson, Brian Scassellati, Mariarosaria Taddeo, Russell Taylor, Manuela Veloso, Zhong Lin Wang, and Robert Wood. The grand challenges of Science Robotics. Science Robotics, 3(14): eaar7650, January 2018. ISSN 2470-9476. doi: 10.1126/scirobotics.aar7650.

18. Thomas B. Sheridan. Human–Robot Interaction: Status and Challenges. Human Factors: The Journal of the Human Factors and Ergonomics Society, 58(4):525–532, June 2016. ISSN 0018-7208, 1547-8181. doi: 10.1177/0018720816644364.

19. Elizabeth Broadbent. Interactions With Robots: The Truths We Reveal About Ourselves. Annual Review of Psychology, 68(1):627–652, January 2017. ISSN 0066-4308, 1545-2085. doi: 10.1146/annurev-psych-010416-043958.

20. John W Krakauer, Asif A Ghazanfar, Alex Gomez-Marin, Malcolm A MacIver, and David Poeppel. Neuroscience needs behavior: correcting a reductionist bias. Neuron, 93(3):480–490, 2017.

21. Steven J Luck and Steven A Hillyard. Spatial filtering during visual search: Evidence from human electrophysiology. Journal of Experimental Psychology: Human Perception and Performance, 20(5):1000, 1994.

22. N. I. Eisenberger. Does Rejection Hurt? An fMRI Study of Social Exclusion. Science, 302(5643):290–292, October 2003. ISSN 0036-8075, 1095-9203. doi: 10.1126/science.1089134.

23. Sylvain Baillet. Magnetoencephalography for brain electrophysiology and imaging. Nature Neuroscience, 20(3):327–339, March 2017. ISSN 1097-6256, 1546-1726. doi: 10.1038/nn.4504.

24. Marco Ferrari and Valentina Quaresima. A brief review on the history of human functional near-infrared spectroscopy (fNIRS) development and fields of application. NeuroImage, 63 (2):921–935, November 2012. ISSN 10538119. doi: 10.1016/j.neuroimage.2012.03.049.

25. Wolfgang H. R. Miltner, Christoph H. Braun, and Michael G. H. Coles. Event-Related Brain Potentials Following Incorrect Feedback in a Time-Estimation Task: Evidence for a “Generic” Neural System for Error Detection. Journal of Cognitive Neuroscience, 9(6):788– 798, November 1997. ISSN 0898-929X, 1530-8898. doi: 10.1162/jocn.1997.9.6.788.

26. Hein T van Schie, Rogier B Mars, Michael G H Coles, and Harold Bekkering. Modulation of activity in medial frontal and motor cortices during error observation. Nature Neuroscience, 7(5):549–554, May 2004. ISSN 1097-6256, 1546-1726. doi: 10.1038/nn1239.

27. Artur Czeszumski, Benedikt V. Ehinger, Basil Wahn, and Peter König. The Social Situation Affects How We Process Feedback About Our Actions. Frontiers in Psychology, 10:361, February 2019. ISSN 1664-1078. doi: 10.3389/fpsyg.2019.00361.

28. Pawel J. Matusz, Suzanne Dikker, Alexander G. Huth, and Catherine Perrodin. Are We Ready for Real-world Neuroscience? Journal of Cognitive Neuroscience, 31(3):327–338, March 2019. ISSN 0898-929X, 1530-8898. doi: 10.1162/jocn_e_01276.

29. Artur Czeszumski, Sara Eustergerling, Anne Lang, David Menrath, Michael Gerstenberger, Susanne Schuberth, Felix Schreiber, Zadkiel Zuluaga Rendon, and Peter König. Hyper-scanning: A Valid Method to Study Neural Inter-brain Underpinnings of Social Interaction. Frontiers in Human Neuroscience, 14:39, February 2020. ISSN 1662-5161. doi: 10.3389/fnhum.2020.00039.

30. Dari Trendafilov, Gerd Schmitz, Tong-Hun Hwang, Alfred O. Effenberg, and Daniel Polani. Tilting Together: An Information-Theoretic Characterization of Behavioral Roles in Rhythmic Dyadic Interaction. Frontiers in Human Neuroscience, 14:185, May 2020. ISSN 1662-5161. doi: 10.3389/fnhum.2020.00185.

31. Arnaud Delorme and Scott Makeig. EEGLAB: An open source toolbox for analysis of single-trial EEG dynamics including independent component analysis. Journal of Neuroscience Methods, 134(1):9–21, March 2004. ISSN 01650270. doi: 10.1016/j.jneumeth.2003.10.009.

32. Andreas Widmann, Erich Schröger, and Burkhard Maess. Digital filter design for electrophysiological data–a practical approach. Journal of neuroscience methods, 250:34–46, 2015.

33. J. A. Palmer, S. Makeig, K. Kreutz-Delgado, and B. D. Rao. Newton method for the ICA mixture model. In 2008 IEEE International Conference on Acoustics, Speech and Signal Processing, pages 1805–1808, Las Vegas, NV, March 2008. IEEE. ISBN 978-1-4244-1483-3. doi: 10.1109/ICASSP.2008.4517982.

34. Olaf Dimigen. Optimizing the ICA-based removal of ocular EEG artifacts from free viewing experiments. NeuroImage, 207:116117, February 2020. ISSN 10538119. doi: 10.1016/j.neuroimage.2019.116117.

35. Luca Pion-Tonachini, Ken Kreutz-Delgado, and Scott Makeig. ICLabel: An automated electroencephalographic independent component classifier, dataset, and website. NeuroImage, 198:181–197, September 2019. ISSN 10538119. doi: 10.1016/j.neuroimage.2019.05.026.

36. Benedikt V. Ehinger and Olaf Dimigen. Unfold: An integrated toolbox for overlap correction, non-linear modeling, and regression-based EEG analysis. PeerJ, 7:e7838, October 2019. ISSN 2167-8359. doi: 10.7717/peerj.7838.

37. GN Wilkinson and CE Rogers. Symbolic description of factorial models for analysis of variance. Journal of the Royal Statistical Society: Series C (Applied Statistics), 22(3):392– 399, 1973.

38. Armand Mensen and Ramin Khatami. Advanced EEG analysis using threshold-free cluster-enhancement and non-parametric statistics. NeuroImage, 67:111–118, February 2013. ISSN 10538119. doi: 10.1016/j.neuroimage.2012.10.027.

39. B. V. Ehinger, P. König, and J. P. Ossandon. Predictions of Visual Content across Eye Movements and Their Modulation by Inferred Information. Journal of Neuroscience, 35(19):7403–7413, May 2015. ISSN 0270-6474, 1529-2401. doi: 10.1523/JNEUROSCI.5114-14.2015.

40. Benedikt V. Ehinger. EEGVIS toolbox, 2018.

41. Yuri G Pavlov, Nika Adamian, Stefan Appelhoff, Mahnaz Arvaneh, Christopher Benwell, Christian Beste, Amy Bland, Daniel E Bradford, Florian Bublatzky, Niko Busch, et al. # eegmanylabs: Investigating the replicability of influential eeg experiments. 2020.

42. Michelle M.E. van Pinxteren, Ruud W.H. Wetzels, Jessica Rüger, Mark Pluymaekers, and Martin Wetzels. Trust in humanoid robots: Implications for services marketing. Journal of Services Marketing, 33(4):507–518, August 2019. ISSN 0887-6045, 0887-6045. doi: 10.1108/JSM-01-2018-0045.

43. Kerstin Sophie Haring, Yoshio Matsumoto, and Katsumi Watanabe. How Do People Perceive and Trust a Lifelike Robot. In Proceedings of the world congress on engineering and computer science, volume 1, page 6, 2013.

44. Munjal Desai, Poornima Kaniarasu, Mikhail Medvedev, Aaron Steinfeld, and Holly Yanco. Impact of robot failures and feedback on real-time trust. In 2013 8th ACM/IEEE International Conference on Human-Robot Interaction (HRI), pages 251–258, Tokyo, Japan, March 2013. IEEE. ISBN 978-1-4673-3101-2 978-1-4673-3099-2 978-1-4673-3100-5. doi: 10.1109/HRI.2013.6483596.

45. Michael Lewis, Katia Sycara, and Phillip Walker. The role of trust in human-robot interaction. In Hussein A. Abbass, Jason Scholz, and Darryn J. Reid, editors, Foundations of Trusted Autonomy, pages 135–159. Springer International Publishing, Cham, 2018. ISBN 978-3-319-64816-3. doi: 10.1007/978-3-319-64816-3âĆĹ.

46. Natalie Sebanz, Günther Knoblich, and Wolfgang Prinz. Representing others’ actions: Just like one’s own? Cognition, 88(3):B11–B21, July 2003. ISSN 00100277. doi: 10.1016/S0010-0277(03)00043-X.

47. J. F. Cavanagh, M. X Cohen, and J. J. B. Allen. Prelude to and Resolution of an Error: EEG Phase Synchrony Reveals Cognitive Control Dynamics during Action Monitoring. Journal of Neuroscience, 29(1):98–105, January 2009. ISSN 0270-6474, 1529-2401. doi: 10.1523/JNEUROSCI.4137-08.2009.

48. Nick Yeung, Matthew M Botvinick, and Jonathan D Cohen. The neural basis of error detection: Conflict monitoring and the error-related negativity. Psychological review, 111(4):931, 2004.

49. Greg Hajcak, Jason S Moser, Clay B Holroyd, and Robert F Simons. The feedback-related negativity reflects the binary evaluation of good versus bad outcomes. Biological psychology, 71(2):148–154, 2006.

50. Nina-Alisa Hinz, Francesca Ciardo, and Agnieszka Wykowska. ERP markers of action planning and outcome monitoring in human – robot interaction. Acta Psychologica, 212:103216, January 2021. ISSN 00016918. doi: 10.1016/j.actpsy.2020.103216.

51. Samuel A. Nastase, Ariel Goldstein, and Uri Hasson. Keep it real: Rethinking the primacy of experimental control in cognitive neuroscience. NeuroImage, 222:117254, November 2020. ISSN 10538119. doi: 10.1016/j.neuroimage.2020.117254.

52. Arianna Curioni, Gunther Knoblich, Natalie Sebanz, A Goswami, and P Vadakkepat. Joint action in humans: A model for human-robot interactions. Humanoid Robotics: A Reference, eds Goswami A, Vadakkepat P (Springer, Dordrecht, The Netherlands), pages 2149–2167, 2019.

53. Basil Wahn and Alan Kingstone. Humans share task load with a computer partner if (they believe that) it acts human-like. Acta Psychologica, 212:103205, January 2021. ISSN 00016918. doi: 10.1016/j.actpsy.2020.103205.

54. Madeline Huberth, Tysen Dauer, Chryssie Nanou, Irán Román, Nick Gang, Wisam Reid, Matthew Wright, and Takako Fujioka. Performance monitoring of self and other in a turn-taking piano duet: A dual-eeg study. Social neuroscience, 14(4):449–461, 2019.

